# Investigations on SARS-CoV-2 and other coronaviruses in mink farms in France at the end of the first year of COVID-19 pandemic

**DOI:** 10.1101/2023.02.02.526749

**Authors:** Marine Wasniewski, Franck Boué, Céline Richomme, Etienne Simon-Lorière, Sylvie Van der Werf, Flora Donati, Vincent Enouf, Yannick Blanchard, Véronique Beven, Estelle Leperchois, Bryce Leterrier, Sandrine Corbet, Meriadeg Le Gouil, Elodie Monchatre-Leroy, Evelyne Picard-Meyer

**Affiliations:** Lyssavirus Unit, Nancy Laboratory for Rabies and Wildlife, ANSES, Malzéville, France; Wildlife Surveillance and Eco-epidemiology Unit, Nancy Laboratory for Rabies and Wildlife, ANSES, Malzéville, France; G5 Evolutionary Genomics of RNA Viruses, Université Paris Cité, Institut Pasteur, Paris, France; Molecular Genetics of RNA Viruses, CNRS UMR 3569, Université Paris Cité, Institut Pasteur, Paris, France; National Reference Center for Respiratory viruses, Université Paris Cité, Institut Pasteur, Paris, France; Mutualized Platform of Microbiology, Pasteur International Bioresources Network, Université Paris Cité, Institut Pasteur, Paris, France; Unit of Viral Genetics and Biosafety, Ploufragan-Plouzané-Niort Laboratory, ANSES, Ploufragan, France; INSERM U1311 DynaMicURe, UNICAEN, UNIROUEN, Normandie University, Caen, France; Virology Department, Caen University Hospital, Caen, France; Nancy Laboratory for Rabies and Wildlife, ANSES, Malzéville, France

**Keywords:** American mink, *Neovison vison*, SARS-CoV-2, *Alphacoronavirus*

## Abstract

Soon after the beginning of the COVID-19 pandemic in early 2020, the *Betacoronavirus* SARS-CoV-2 infection of several mink farms breeding American minks (*Neovison vison*) for fur was detected in several countries of Europe. The risk of a new reservoir formation and of a reverse zoonosis from minks was then a major concern. The aim of this study was to investigate the four French mink farms for the circulation of SARS-CoV-2 at the end of 2020. The investigations took place during the slaughtering period thus facilitating different types of sampling (swabs and blood). In one of the four mink farms, 96.6% of serum samples were positive in SARS-CoV-2 ELISA coated with purified N protein recombinant antigen and 54 out of 162 (33%) pharyngo-tracheal swabs were positive by RT-qPCR. The genetic variability among 12 SARS-CoV-2 genomes sequenced in this farm indicated the co-circulation of several lineages at the time of sampling. All SARS-CoV-2 genomes detected were nested within the 20A clade (Nextclade), together with SARS-CoV-2 genomes from humans sampled at the same period. The percentage of SARS-CoV-2 seropositivity by ELISA varied between 0.5 and 1.2% in the three other farms. Interestingly, among these three farms, 11 pharyngo-tracheal swabs and 3 fecal pools from two farms were positive by end-point RT-PCR for an *Alphacoronavirus* highly similar to a mink coronavirus sequence observed in Danish farms in 2015. In addition, a mink *Caliciviridae* was identified in one of the two positive farms for *Alphacoronavirus*. The clinical impact of these unapparent viral infections is not known. The co-infection of SARS-CoV-2 with other viruses in mink farms could contribute to explain the diversity of clinical symptoms noted in different infected farms in Europe. In addition, the co-circulation of an *Alphacoronavirus* and SARS-CoV-2 within a mink farm would increase potentially the risk of viral recombination between alpha and betacoronaviruses already suggested in wild and domestic animals, as well as in humans.

**Author summary:** France is not a country of major mink fur production. Following the SARS-CoV-2 contamination of mink farms in Denmark and the Netherlands, the question arose for the four French farms.

The investigation conducted at the same time in the four farms revealed the contamination of one of them by a variant different from the one circulating at the same time in Denmark and the Netherlands mink farms.

Investigation of three other farms free of SARS-CoV-2 contamination revealed the circulation of other viruses including a mink Alphacoronavirus and *Caliciviridae*, which could modify the symptomatology of SARS-CoV-2 infection in minks.

## 1. INTRODUCTION

Soon after the beginning of the COVID-19 pandemic in early 2020, the SARS-CoV-2 infection of several mink farms breeding American minks (*Neovison vison*) for fur, was detected in Europe, in the Netherlands first (1), in April 2020, and then in Denmark (2). Infections with SARS-CoV-2 in mink farms were then detected in several other countries in Europe: in Italy, Spain, Poland, Greece, Lithuania and Sweden (3). In Denmark, SARS-CoV-2 has spread rapidly in each farm and among mink farms, and was associated with the emergence of a specific variant (called cluster 5) detected in November 2020. At the time, this SARS-CoV-2 variant was believed to present different phenotypic characteristics including escape from neutralizing antibodies (4,5). Moreover, if the primary contamination of the mink farms was due to human infection (6), back transmission from minks to humans was detected in the Netherlands (7) and Poland (8).

In early 2020, four American mink farms of limited size (less than 1000 to about 15,000 minks (according to the French Ministry of Agriculture) were in operation in France. These farms were located in different regions. No clinical sign had been observed in minks from these farms since the beginning of the pandemic. The aim of the present study was to investigate the circulation of SARS-CoV-2 and potentially other coronaviruses in the four French mink farms. As mink farming has a seasonal production, with slaughtering of the young adults of the year at the end of the same year, the investigations took place at the end of 2020 to conduct the most exhaustive survey as possible on the individuals born in 2020. Different types of samples were collected during the slaughtering period. Serology and viral RNA detection in the upper and lower respiratory tract and in feces were performed as well as Bayesian analysis to determine the potential circulation of different lineages in French mink farms.

## 2. RESULTS

### 2.1. Sample collection

The four American mink farms present in France (named A to D) were investigated (Figure 1).

**Fig. 1:**
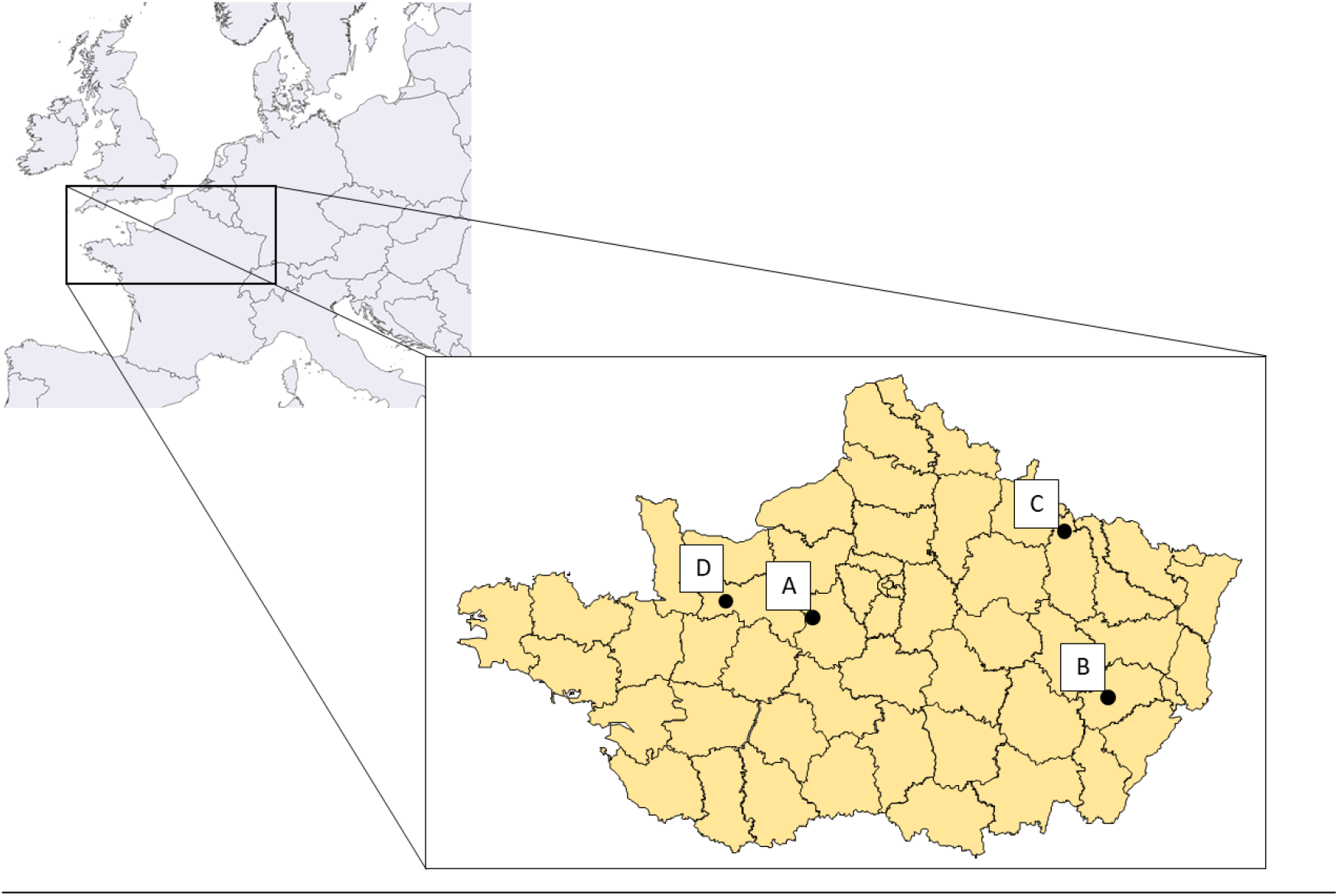
Location of the American mink farms in France in November 2020. (A) Eure-et-Loir, (B) Haute-Saône, (C) Meuse and (D) Orne.

A total of 1912 minks born in 2020 were sampled for blood, with a minimum of 60 animals per building, and 1643 pharyngo-tracheal swabs were collected. Characteristics of the farms and of the sampling are presented in Table 1.

**Table 1:** Location (department) of mink farms, number of blood samples by route of sampling, swab samples and fecal samples collected in November 2020 in France on adult minks of the year

| Farm | Minks present in the farms | Nb. of scanstars | Nb. scanstars sampled from housings with slaughtered minks for fur | Blood samples |  | Pharyngo--tracheal swabs | Nb. Scanstars sampled from housing with unslaughtered minks for breeding | Fecal samples (nb. of pools collected in each scanstars) |
| --- | --- | --- | --- | --- | --- | --- | --- | --- |
|  |  |  |  | Intra-cardiac puncture | Retro-orbital |  |  |  |
| A | 3,800 | 3 | 3 | 179 |  | 162 | 0 | 0 |
| B | 950 | 2 | 2 | 120 |  | 120 | 0 | 0 |
| C | 7,900 | 15 | 9 |  | 403 | 395 | 6 | 56*<sup>1</sup> |
| D | 10,850 | 26 | 20 | 1210 |  | 1230 | 6 | 34*<sup>2</sup> |
| Total | 23, 500 | 47 | 34 | 1912 |  | 1643 | 13 | 90 |

\*<sup>1</sup> number of fecal samples collected / number of minks sampled: 1-10 / 2-5 animals
\*<sup>2</sup> number of fecal samples collected / number of minks sampled: 4-7 / 10 animals

### 2.2. Anti-SARS-CoV-2 antibodies detection (ELISA and seroneutralization test)

All mink blood samples were tested by a SARS-CoV-2 ELISA coated with purified N protein recombinant antigen. Among them, samples presenting a doubtful or positive result were systematically tested again by seroneutralization assay to confirm or overturn the ELISA results.

For farm A, where evidence of ongoing infection was noted (SARS-CoV-2 RNA was found in pharyngo-tracheal swabs), we randomly selected 16 samples among the positive sera to be tested by seroneutralization assay. For farms B, C and D, samples that were negative by ELISA were also tested by seroneutralization assay as negative controls.

The results of the ELISA are shown in Table 2. The percentage of seropositivity varied between 0.5 and 1.2% in non-infected farms and reached 96.6% in the infected farm (A). For farm A, the S/P% values of positive samples ranged from 62.3 to 492.6 with mean value equal to 307.9 (N=173).

**Table 2:** Serological results (positive/doubtful/negative) obtained by ELISA on mink serum samples collected in the four mink farms in France in November 2020

| Farm | Samples | Positive | Doubtful | Negative | % of positive (including doubtful results) |
| --- | --- | --- | --- | --- | --- |
| A | 179 | 173 | 1 | 5 | 96.6 |
| B | 120 | 1 | 0 | 119 | 0.8 |
| C | 403 | 1 | 1 | 401 | 0.5 |
| D | 1210 | 13 | 1 | 1196 | 1.2 |

The samples found positive or doubtful from farms B, C and D, as well as a few randomly selected positive samples from farm A, were tested by seroneutralization assay (Table 3).

**Table 3:**
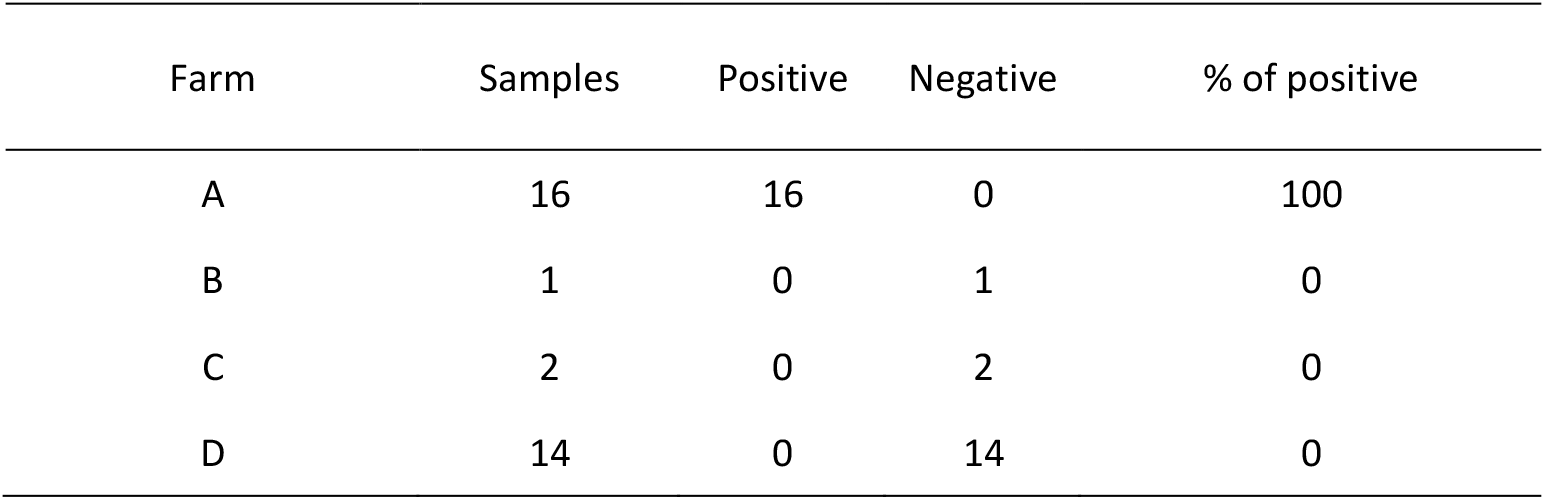
Serological results obtained by using the seroneutralization assay on ELISA positive/doubtful mink serum samples collected in the four mink farms in France in November 2020

For the three non-infected farms, no SARS-CoV-2 neutralizing antibody was detected by seroneutralization assay in the positive or doubtful sera obtained by ELISA test (Table 3). However, for farm A, SARS-CoV-2 neutralizing antibodies were detected in the 16 ELISA positive samples. Neutralizing titers ranged from 81 till above 2239, confirming the circulation of the virus in this farm.

### 2.3 Detection of SARS-CoV-2 viral RNA in pharyngo-tracheal swabs

The TaqMan RT-qPCR analysis of pharyngo-tracheal swabs revealed that farm A was experiencing an ongoing SARS-CoV-2 outbreak: of the 162 minks tested from farm A, SARS-CoV-2 RNA was detected in 54 swabs at the date of sampling (Table 4). Of the 54 positive samples, 33 of them showed low levels of viral RNA (Ct value > 32, which corresponds to approximately 0-5copies/μL of RNA). The 21 positive minks for SARS-CoV-2 RNA (Ct values ranging from 18.8 to 31) were found positive with RNA titers ranging from 10 to 3,79E+04 copies/μL of RNA.

**Table 4:**
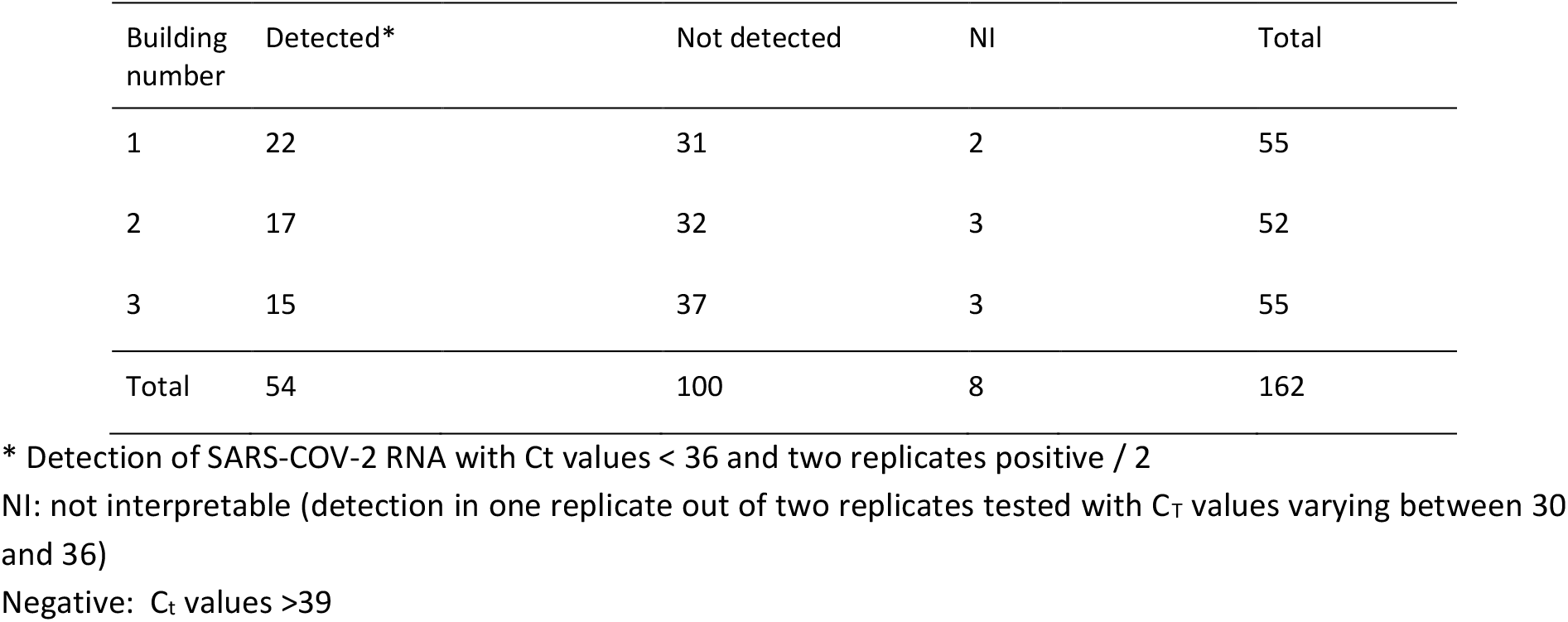
SARS-CoV-2 RNA detection in pharyngo-tracheal swabs in farm A.

| Building number | Detected* | Not detected | NI | Total |
| --- | --- | --- | --- | --- |
| 1 | 22 | 31 | 2 | 55 |
| 2 | 17 | 32 | 3 | 52 |
| 3 | 15 | 37 | 3 | 55 |
| Total | 54 | 100 | 8 | 162 |

\* Detection of SARS-COV-2 RNA with Ct values < 36 and two replicates positive / 2
NI: not interpretable (detection in one replicate out of two replicates tested with Ct values varying between 30 and 36)
Negative: Ct values >39

No SARS-CoV-2 RNA was detected in samples from the three other farms (n= 1481 samples tested).

### 2.4. SARS-CoV-2 genomes

SARS-COV-2 RNA samples with Ct values less than 31 (12.5-3.1E+02 copies/μL of RNA) were submitted to full genome analysis. Nearly complete viral genome sequences (coverage > 99.4%, average depth > 15 000, maximum depth > 100 000) were obtained from each of the 12 nucleic acid extracts positive for SARS-CoV-2 by RT-qPCR (Ct values of 26.2-30). An additional extract, weakly positive by RT-qPCR, yielded partial sequences corresponding to different regions of the genome. All twelve genomes (GISAID under numbers EPI_ISL_1392906 & EPI_ISL_10036487-97) were classified as 20A based on a panel of mutations compared to reference sequence NC_045512 and a Nextclade analysis (Fig. S1, Table S1). A Bayesian phylogenetic analysis using all high-quality mink derived SARS-CoV-2 genomes available in the GISAID database clustered SARS-CoV-2 from the mink farm in France together in a monophyletic clade supported by significant node value (Posterior probability) and a relatively long branch (Fig. 2). This clear clustering does not link the SARS-CoV-2 from the mink farm in France to any another mink farm.

**Figure 2.**
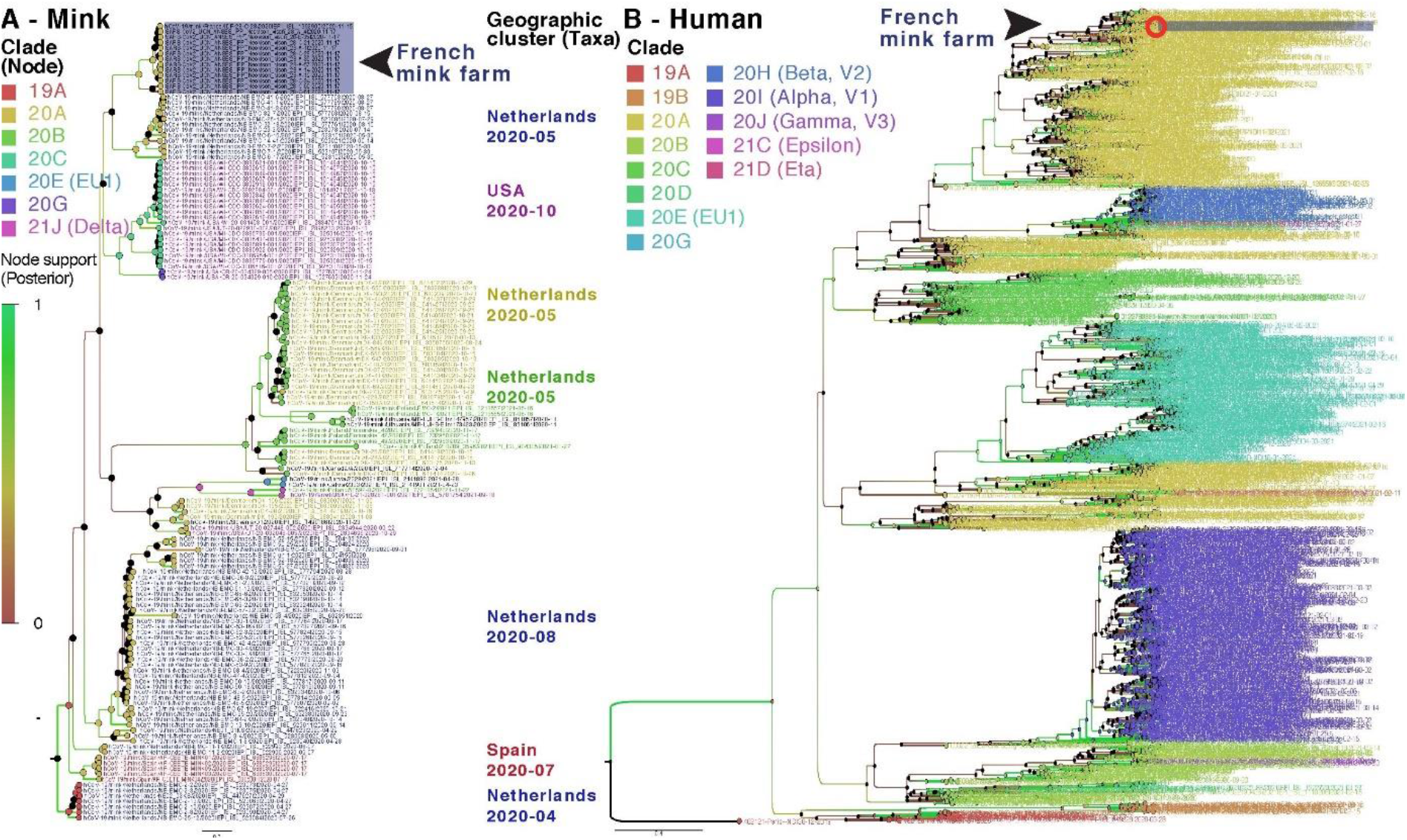
Bayesian phylogenies on a mix of representative set of SARS-CoV-2 sequences available in GISAID. SARS-CoV-2 sequences detected in the mink farm in France are highlighted in purple. **A. Bayesian phylogeny of all SARS-CoV-2 genomes from minks over the study period**. Node and taxa labels are colored according to nextstrain (https://nextstrain.org) clade classification and main geographic clusters, respectively. **B. Bayesian phylogeny of human derived SARS-CoV-2 genomes collected in France and mink derived SARS-CoV-2 genomes described in this study**. A diversity optimized dataset was obtained by collecting and filtering complete, high quality genomes, of SARS-CoV-2 detected in France (of Human and mink origin). Taxa are colored according to nextstrain clades classification.

Moreover, thirteen SNPs (Single Nucleotide Polymorphisms) were specific to this clade by contrast to other SARS-CoV-2 clades detected in mink (*Neovison vison*) worldwide at the time of data collection and phylogenetic analyses based on 821 mink SARS-CoV-2 genomes, retrieved on January 5, 2022 (Fig. 3; Table S1). Among these SNPs, differentiating mink SARS-CoV-2 sequences from France from other available mink SARS-CoV-2 sequences, 4 induce an amino-acid change (V676L, K1141R, E1184D in the Orf1b and S477N in the Spike). Within this monophyletic clade, genetic variability was observed among SARS-CoV-2 genomes from the French farm (including non-silent mutations), indicating the co-circulation of several variants in this setting at the time of sampling (Tables S1 and S2). A Bayesian phylogenetic analysis using the 2020’s year of data from GISAID (437 human SARS-CoV-2 in a diversity optimized matrix based on 99,875 % pairwise nucleotide identity cutoff from 42700 genomes) showed that SARS-CoV-2 genomes detected in the mink farm in France nested within the 20A clade, together with SARS-CoV-2 genomes from humans sampled at the same period (Fig. S2). In addition, the non-silent mutation in S (S477N) that differentiates these mink SARS-CoV-2 genomes from that of other SARS-CoV-2 sampled in mink elsewhere, was also observed in SARS-CoV-2 sampled in humans in France at the same period (Table S3 and Fig. S2).

**Figure 3.**
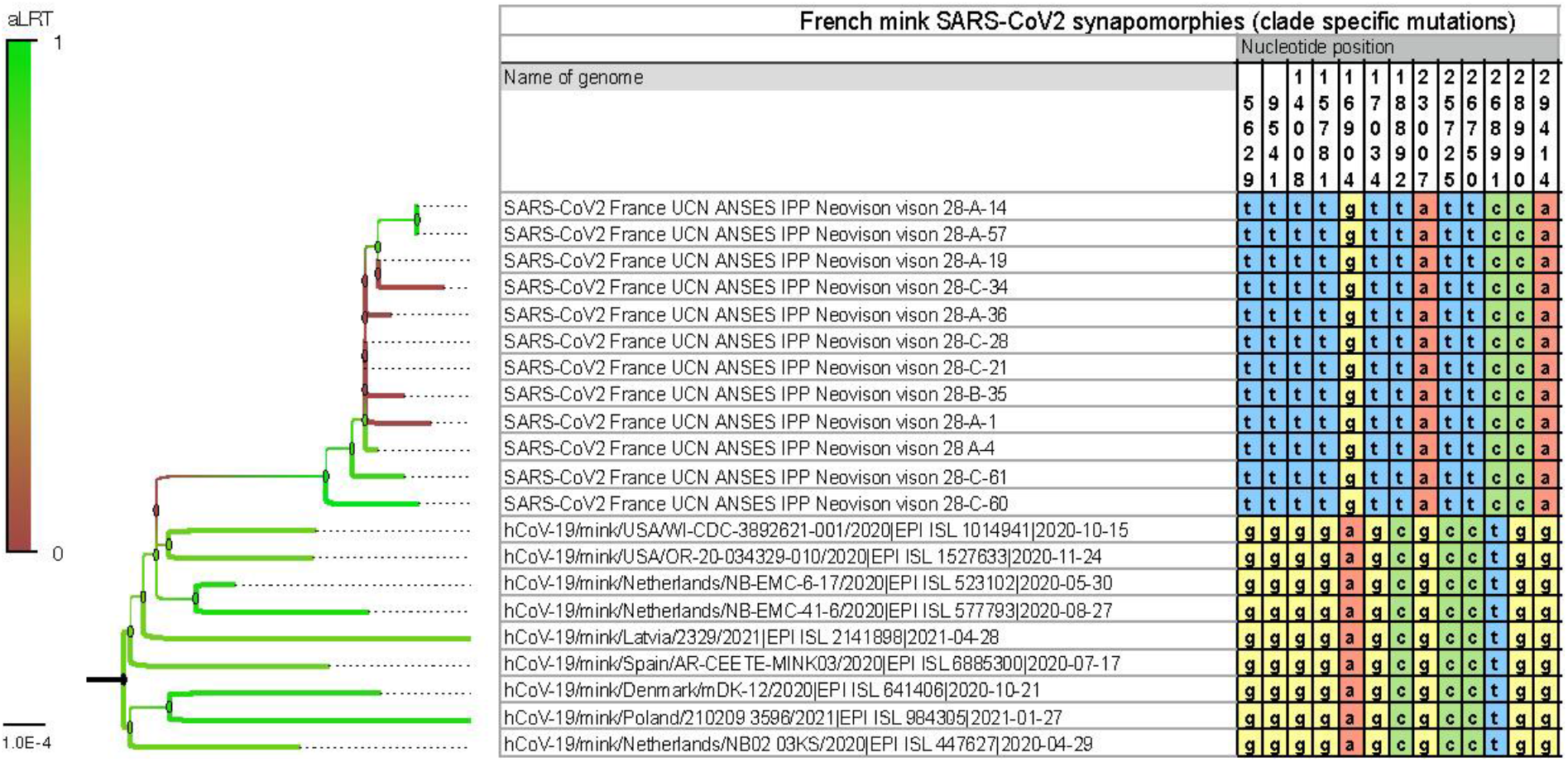
SARS-CoV-2 synapomorphies in American minks from a mink farm in France with the thirteen SNPs specific to the clade formed by the mink SARS-CoV-2 in France compared to other mink SARS-CoV-2 located elsewhere. SNPs are represented with the nucleotide positions on the SARS-CoV-2 genome, and in yellow for G, red for A, green for C and blue for T. Branchs are colored according to the bootstrap values.

### 2.5. Detection of an *Alphacoronavirus* in pharyngo-tracheal swabs and feces

We analyzed the pharyngo-tracheal swabs and feces for the presence of RNA coronaviruses by end-point RT-PCR targeting the pol gene. The end-point RT-PCR analysis of pharyngo-tracheal swabs revealed that two out of three farms were positive for RNA coronaviruses distinct from SARS-CoV-2. Of 236 swab samples tested, 11 minks were positive for *Alphacoronavirus* RNA, with respectively two samples in farm C and nine in farm D (Table 5). No coronavirus was detected in swabs from farm B. Farm A samples were not analyzed for the presence of the A*lphacoronavirus* genome. A total of 90 feces pools (56 from farm C and 34 from farm D) were simultaneously tested by RT-PCR targeting the pol gene and by beta-actin RT-PCR for investigating the presence of RNA inhibitors. The beta-actin housekeeping gene RNA was not detected in 46.6% of the fecal samples tested (n=42/90*100) with the highest proportion of not exploitable samples in farm C (55.4%=31/56*100) compared to farm D (32%=11/34*100). Of 48 exploitable feces pools, 3 were positive for *Alphacoronavirus* RNA: one in farm C and 2 in farm D.

**Table 5:**
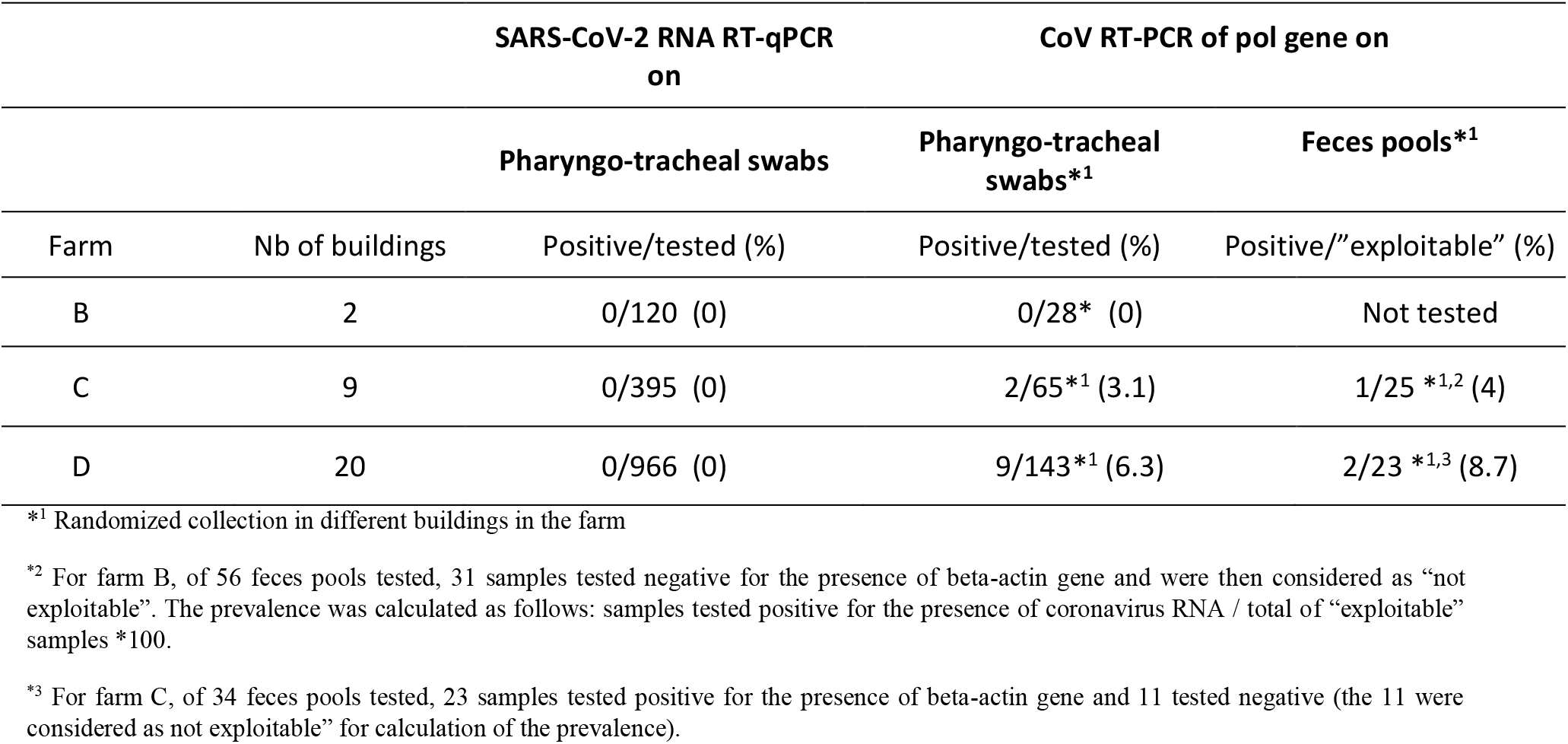
Detection of α-CoV in pharyngo-tracheal swabs and feces in negative SARS-CoV-2 mink farms in France

Of the 14 samples tested positive by RT-PCR for the pol gene, 5 were weakly positive and then were either not submitted or failed the SANGER sequencing. Sequence analyses of PCR products (n=9) showed that all sequenced samples (7 pharyngo-tracheal swabs and 2 fecal pools) by SANGER were alphacoronaviruses. Of the 9 nucleotide consensus sequences, 8 were included in the phylogenetic analysis, all of them grouped within *Alphacoronavirus, Minacovirus* subgenus (bootstrap=95), close to the mink coronavirus (anciently named ferret coronavirus - Figure 4). The ninth sample 61-B23-2 was not included in the phylogeny, due to the low quality of the nucleotide sequence obtained by SANGER sequencing and the fact that we were unable to obtain the consensus sequence for this sample. BLAST analysis of the forward sequence still showed that the sample 61-B23-2 is closed to both mink coronavirus 1 MN535737 isolated in Denmark in 2015 and an *Alphacoronavirus* isolated on a mink in China in 2016 (MF113046). High nucleotide similarities were observed for the Pol gene sequences from farm C (n=1) and farm D (n=8). A BLAST search showed that the nine strains shared between 94.6% to 96.3 % identity at the nucleotide level with the mink coronavirus 1 (MN535737) strain isolated in Denmark in 2015.

**Figure 4.**
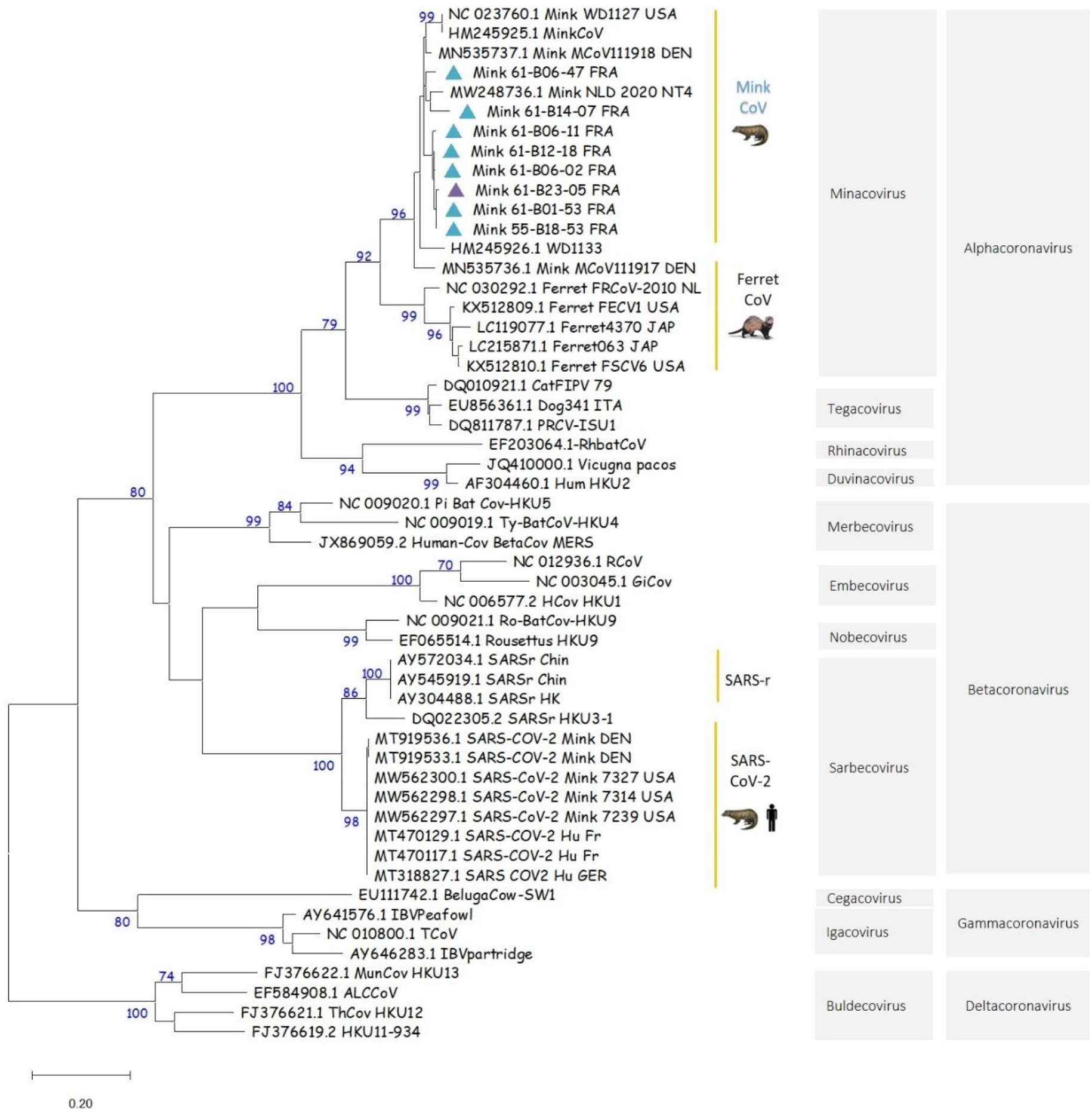
Maximum likelihood (ML) phylogeny inferred with 8 mink coronavirus consensus sequences from France and 45 representative GenBank sequences including alphacoronaviruses (n=25), betacoronaviruses (n=20) with mink and human SARS-CoV-2 sequences, gammacoronaviruses (n=4) and deltacoronaviruses (n=4). Bootstrap values above 70% were considered as statistically significant. Pharyngo-tracheal swabs and feces infected by the mink coronavirus are represented on the tree with a turquoise and purple triangle, respectively. The 8 partial pol gene consensus sequences included in the ML tree are accessible in GenBank under accession numbers: Mink_61-B14-07_FRA: ON985270; Mink_61-B12-18_FRA: ON985271; Mink_61-B06-47_FRA: ON985272; Mink_61-B06-11_FRA: ON985273; Mink_61-B06-02_FRA: ON985274; Mink_61-B01-53_FRA: ON985275; Mink_55-B18-53_FRA: ON985276; Mink_61-B23-05_FRA: ON985277

Of the three fecal samples tested positive for the coronavirus pol gene, the sample (61-B23-05) the most positive was submitted to high throughput sequencing (HTS). The 61-B23-05 extract yielded many gaps throughout the genome, with partial sequences (i.e. 706, 639, 597, 445, and 206 nucleotides) corresponding to five different regions of a Mink strain coronavirus 1 MCoV1/11918-1/DK/2015 (MN535737.1). BLAST analysis performed on the five partial sequences showed 89-95% identity with MN535737.1.

### 2.6. Detection of Mink *Caliciviridae* RNA genome in pharyngeo-tracheal samples

Deep sequencing of viral RNA extracted from pharyngeo-tracheal swab samples 61-B06-11 and 61-B12-04 mainly generated bacteria or host (Mustelidae)-originating reads; only 0.06% to 0.7% of the reads were identified as viral sequences with, for sample 61-B12-04, 75% of the reads identified as mink *Caliciviridae*. None of the reads obtained for these two samples corresponded to *Alphacoronavirus* sequences. After the Spades assembly of the 61-B12-04 reads, one contig (4320 nucleotide long) was identified as a *Caliciviridae* sequence with 95% identity with the *Caliciviridae* strain [Mink/China/2/2016](https://blast.ncbi.nlm.nih.gov/Blast.cgi#_blank) (MF67785). Several reads were identified as mink *Caliciviridae* in the two samples. The whole genome size for the two samples was of 8,427 nt in length and consisted of three open reading frames and two untranslated regions (5’ and 3’) of 13 and 103 nt of length, respectively. ORF1 ranged from nt 14 to nt 5851 (encoding a polyprotein of 1,946 aa), ORF2 ranged from nt 5857 to nt 7899 (681 aa), ORF3 ranged from nt 8130 to nt 8306 (59 aa). The alignment of the truncated sequence 61-B12-04 (GenBank Number OP485683) with the MF677852 mink *Caliciviridae* strain from China showed an amino-acid identity of 99.1% along the whole genome), with 99.4 % for ORF1, followed by 98.4% and 98.3 for ORF2 and ORF3, respectively. Finally, BLAST analysis of the 61-B06-11 and 61-B12-04 sequences showed > 94.4% (E value=0.0; Score=8167; number of hits =5; 4982/5279*100) and 96% (E value= 4e-103; Score= 392; number of hits=5; 230/239*100) of nucleotide identity with the Mink *Caliciviridae* strain from China MF677852, respectively.

## 3. DISCUSSION

This study describes the investigations on SARS-CoV-2 in the four mink farms in operation in France at the end of the first year of the COVID-19 pandemic. The small number of farms and animals per farm made possible to implement a robust transversal study allowing the detection of a minimum prevalence of 5%. SARS-CoV-2 infected minks in one farm, with a high serological prevalence (above 96%) and SARS-CoV-2 RNA was detected in 30% of the sampled minks. Infection had spread in the three buildings of the farm. No clinical sign nor suspicious mortality was observed by the breeder suggesting a mild outbreak which could have been easily missed in the absence of investigations (2,9,10).

All twelve SARS-CoV-2 genome sequences obtained, despite some variability, belonged to the 20A clade, according to Nextclade and two Bayesian phylogenetic reconstructions. Given the genetic relationship between the SARS-CoV-2 sequenced in the mink farm in France and the SARS-CoV-2 responsible for the concomitant epidemic wave of 20A viruses in humans, as well as the previous description of the zoonotic circulation of SARS-CoV-2 between human and minks in the Netherlands, reverse zoonosis appears as the main hypothesis to explain the SARS-CoV-2 circulation detected in the minks of this farm. In addition, comparison with other SARS-CoV-2 sequenced in minks worldwide clearly shows the monophyly of this mink clade found in France. The significant genetic distance to viruses found in other countries is supported by 13 mutations specific to this clade. These analyses reinforce the hypothesis of a local transmission from human to explain the origin of the circulation in minks in the present study. Among these mink clade specific mutations, four were non-silent and one, resulting in the S477N change in the spike, was also present in several genomes sequenced from humans at the same period in France. Despite substantial genetic variability observed here in mink SARS-CoV-2, data was not discriminant enough to test with confidence the monophyly of this mink clade when analyzed with all available human SARS-CoV-2 from the region and sampled at the same period. Bayesian reconstruction shows apparent paraphyly of the mink clade found in France, with some internal human SARS-CoV-2 sub-clustering, but these nodes are not significant (low posterior probabilities). The dataset does not allow to clearly establish whether one or several reverse-zoonotic events occurred. However, the relative genetic identity of genomes detected in the farm is in favor of a contamination of the farm relatively shortly before the sampling, with a viral circulation probably shorter than 3 months. Indeed, this duration is insufficient to allow the diversification in several clearly distinct lineages as the accumulation is around two mutations a month on average (11). A high number of minks can be infected within a short time: Hammer et al. (9) described an increase of prevalence from 4% to 97% in eight days in minks in one farm in Denmark. The detection of SARS-CoV-2 RNA in pharyngo-tracheal swabs with Ct values varying between 18.8 and 38.4 was the sign that infection was recent in a few minks, especially for minks with low Ct values. Indeed, after experimental infection, viral RNA can be detected in the upper respiratory tract from 2 dpi to 17 dpi of exposed minks(12). As the detection of SARS-CoV-2 in farm A induced immediate control measures (culling of all minks), it was not possible to follow further the genomic evolution of the virus in minks and to test the eventuality of spill-back to humans.

No SARS-CoV-2 contamination was detected in the three other farms, but less than 1% of the samples in farms B and C, and 1% in farm D were positive by ELISA, despite a negative seroneutralization test. The specificity of the ELISA test in minks is very high, close to 99%, better than in humans (where the specificity is evaluated between 92.5 and 98.8%) (13,14). In farms C and D, we detected an *Alphacoronavirus*, Mink-coronavirus sequence, found in both pharyngo-tracheal swabs and feces. Mink coronavirus infection is associated with epizootic gastroenteritis (ECG) in minks when in combination with several enteric viruses, but it can also be asymptomatic (15). As ECG is an economic concern, several authors in different countries studied the disease characteristics (distribution, lesions…) (15,16). Then, by considering the ECG frequency found in these studies, mink coronavirus infection in fur minks appears to be not so rare, but to our knowledge, little data exist on prevalence. However, the potential circulation of an *Alphacoronavirus* at the same time as a very active SARS-CoV-2 outbreak within a farm would increase considerably the risk of viral recombination. Recombination of alpha and betacoronaviruses has been already described in wild and domestic animals as well as in humans (17–19). Such an event could result in an evolutionary jump and may generate a recombinant with unpredictable phenotype and fitness that may promote the emergence of a novel coronavirus. This potential issue should be seriously considered in countries where mink farms (or farm breeding of other small carnivores) are insufficiently monitored and where preventive culling of SARS-CoV-2 positive farms is not applied.

Finally, by attempting to obtain the complete genome sequence of mink coronavirus on swab samples that were positive for *Alphacoronavirus* by end-point RT-PCR, a nearly full genome of a mink *Caliciviridae* was obtained by HTS. The *Caliciviridae* sequence was predominant in our two samples in comparison with that of the mink coronavirus for which the sequence have not been obtained. The absence of detection of mink coronavirus by HTS was disappointing but is linked to the very low load of coronavirus RNA, which was only detected after a nested PCR. As a result, coronavirus reads were hidden by the high proportion of bacteria and host (mustelidae) reads accounting for ∼99% of the reads in this sample. HTS sensitivity is known to be much lower than that of PCR, even more so with nested PCR. Similar *Caliciviridae* have already been sequenced in minks in the USA (20) and China (21). The associated clinical signs are not completely identified; some authors described no clinical sign (22) or diarrhea (22) or hemorrhagic pneumonia (20). Beside the results obtained on coronaviruses, our study on the French mink farms also shows that a number of potentially pathogenic agents were silently circulating. This also seems to be the case in other farms (22). These infections could modify the symptomatology of SARS-CoV-2 infection in minks, which has shown more or less acute clinical signs depending on the farms in the different affected countries.

## 4. MATERIALS AND METHODS

The four American mink farms present in France (named A to D) were located in rural areas isolated from human habitations. They were small family farms managed by one person, except farm D where external workers were regularly employed. In each farm, minks were housed in wire netting cages placed in scanstars, each of this housekeeping unwalled building consisting of two rows of several dozen cages. A surrounding fence assured protection from intrusion in the farms. During the investigations in November 2020, each cage housed two adult minks. Each cage was equipped with automatic water distribution, and food was distributed every two days. According to the farmers’ declaration, no material sharing between farms and no animal exchange has been recorded.

Before slaughtering, the four mink populations consisted of 3,800 minks in farm A, 950 in farm B, 7,900 in farm C and 10,850 in farm D, spread in 3, 2, 15 and 26 housekeeping buildings respectively.

### 4.1. Mink sampling and samples collection

Each scanstar was considered as an independent epidemiological unit. Sixty animals were sampled in each to enable sensitive detection, assuming a minimum apparent prevalence of 5%, with a 95% confidence interval.

Minks were sampled during the slaughtering period (10 to 26 November 2020). Blood was sampled on freshly euthanized mink carcasses, by intra-cardiac puncture in farms where carving up was realized in place (farms A, B and D) or by retro-orbital sampling in farm C where animals were stored intact before fur treatment. Blood samples were stored at 4°C before being transferred to the laboratory. Samples were allowed to clot and then centrifuged (1000g, 15 min) to obtain serum. Sera were stored at -20°C until serological testing. Sera were heat-inactivated at 56°C during 30 min preceding seroneutralization assay.

Pharyngo-tracheal swabs were collected on mink carcasses by tracheal retro-route in farms A, B and D, and by oropharyngeal route in farm C. The swabs were immediately kept in a volume of 500 μL of cell culture medium (DMEM) supplemented with 1% of antibiotics (mix of Penicillin, Streptomycin and Amphotericin B) before being frozen in liquid nitrogen to ensure viral integrity. The samples were transferred to the laboratory and stored at < -70°C until RNA extraction.

The sample collection was completed with feces samples of breeding animals in farms C and D. Indeed, in these two farms, some scanstars housed the unslaughtered animals that would serve as breeding stock. Blood samples and tracheal swabs were then impossible to achieve. Pools of feces were collected in farms C and D in 6 individual scanstars, with a total of 56 and 34 pool samples for the two farms, respectively. Samples were stored at < -70°C until RNA extraction.

### 4.2. SARS-CoV-2 isolate

SARS-CoV-2 strain UCN19 was amplified on cells as described previously (23) and used at passage 2 for the seroneutralization assays.

### 4.3. Enzyme Linked Immunosorbent Assay (ELISA) test

The Enzyme Linked Immunosorbent Assay provided by IDVet (ID Screen®ELISA, SARS-CoV-2 Double Antigen Multi-species) has been used as a screening tool to detect the antibody presence in mink sera. This ELISA is a double antigen ELISA for the detection of antibodies directed against the nucleocapsid of SARS-CoV-2 in animal serum, plasma or whole blood. The samples were tested according to the manufacturer’s recommendations and as previously described (24,25). Briefly, 25 μL of each serum sample was added in the microplate and diluted 1:2 in sample diluent. Microplates were incubated 45 minutes at 37 °C+/-2°C. Five washings were performed after incubation. Then, 100 μL of the conjugate (a purified recombinant N protein antigen labeled with horseradish peroxidase) were distributed to each well. The microplates were incubated for 30 min at 21°C +/-5°C. Five washings were performed to remove the unbound conjugate. The presence of the complex antibodies/conjugate was revealed by adding 100μL of TMB (TetraMethylBenzidine) chromogen solution to each well. The microplates were incubated in the dark for 20 min at 21 °C +/-5°C. The enzymatic reaction was stopped by adding 100μl of a stop solution. The microplates were read at 450 nm. Positive and negative controls provided by the manufacturer were used to validate each test plate.

The conditions of validation described by the manufacturer were implemented to validate the tests and to interpret the results obtained for the different samples. The “Sample/Positive” ratio was calculated as follows and expressed as a percentage (S/P%) :

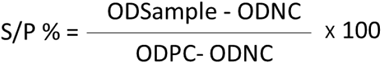

where ODNC is the optical density of the negative control and ODPC is the optical density of the positive control and ODSample is the optical density of the sample. According to cut-off determined by the manufacturer, three kind of results could be obtained:

- If S/P% is below or equal to 50%, the sample was considered negative
- If S/P% is between 50% and 60%, the sample was considered doubtful
- If S/P% is above or equal to 60%, the sample was considered positive

### 4.4. Seroneutralization assay

Briefly, in 96-well microplates, 200μL of VERO E6 cell suspensions in Dulbecco’s Modified Eagle medium (DMEM) containing 10% FCS (Fetal calf serum) and 1% antibiotics (Penicillin/Streptomycin) were added to each well, representing 20 000 cells per well, 24 hours before starting the seroneutralization assay.

Each serum sample as well as positive and negative internal controls were distributed in two consecutive wells of 96-well microplates, and then serially diluted with a dilution step of 1 to 3 within DMEM containing 10% FCS and 1% antibiotics (Penicillin/Streptomycin). Then 50μL of SARS-CoV-2 virus diluted in medium containing around a 50% tissue culture infective dose (TCID50) of 100 per 50μL (checked by back-titration during the seroneutralization assays) were added to each well containing samples and internal controls. The plates were incubated at 37°C/5% CO2 for 1h to allow neutralization complexes to be formed between the neutralizing antibodies and the virus. At the end of the incubation, the supernatant fluid was removed from each well of the plates containing VERO E6 cell suspensions and immediately 100μL of the mix of the virus and serially diluted samples or controls were transferred to individual wells on the cell layer. The microplates were incubated at 37°C in a humid chamber containing 5% CO2, at least 3 days post-infection. Then, plates were qualitatively read according to an “all or nothing” scoring method for the presence of viral cytopathic effect (CPE). The neutralization titers were assigned to each serum based on the highest dilution that prevented discernible cytopathic effect.

### 4.5. RNA extraction

#### Pharyngo-tracheal swabs

Viral RNA was extracted from 140 μL of pharyngo-tracheal swabs medium. Viral RNA extraction was performed by using the Qiagen Viral RNA mini kit according to the manufacturer’s instructions (Qiagen, Les Ulis, France), with minor modifications. To inactivate potential infectious status of samples by SARS-CoV-2, a volume of 15 μL of Triton X-100 (MP Biomedicals, Illkirch, France) was added to 560 μL of AVL Lysis buffer (Qiagen, Courtaboeuf, France) for each sample testing. RNA was eluted in a final volume of 60 μL and stored at < -70°C. A negative RNA extraction control was performed for each set of 24 samples tested.

#### Fecal samples

100 mg of fecal sample were placed in a lysing matrix E tube containing beads (MP Biomedicals Germany Gmbh, Eschwege, France) and filled with 1ml of CTAB buffer (Promega France, Charbonnières les Bains, France). Tubes were placed in a shaking heat block at 65°C, vigorously vortexed for 1 min and mixed with 40 μL of Proteinase K Solution (Promega France, Charbonnières les Bains, France). After an incubation at 70°C for 10 min, the lysates were grinded for 30 sec at 7.0 m/s six times in a Fast Prep-24 ™ 5G bead beater (MP Biomedicals Germany Gmbh, Eschwege, France) then centrifuged at 10,000 x g for 5 min. A volume of 300 μL of clear lysate was transferred to a tube containing 300 μL of Lysis buffer. RNA extraction was performed with the Maxwell RSC PureFood GMO and Authentication kit using a Maxwell RSC instrument (Promega France, Charbonnières les Bains, France), according to the manufacturer’s instructions. RNA was eluted in a final volume of 100μL and stored at < -70°C.

Negative (non-template control) and positive (hedgehog betacoronavirus) controls (RNA extraction control) were performed for each set of 16 samples tested.

### 4.6. TaqMan RT-qPCR of E gene using specific SARS-CoV-2 primers

TaqMan RT-qPCR was performed as previously described (23). Coronavirus primers (E_Sarbeco_F (forward): 5’-ACAGGTACGTTAATAGTTAATAGCGT and E_Sarbeco_R (reverse): 5’-ATATTGCAGCAGTACGCACACA) and probe (E_Sarbeco_P1: 5’-FAM-ACACTAGCCATCCTTA CTGCGCTTCG-BHQ-1) targeting the envelope protein gene (E gene) were used for the study (26). Primers and probe were provided by Eurogentec (Angers, France). TaqMan RT-qPCR assays were performed in a total volume of 25 μL containing 2.5 μL of RNA sample, 12.5 μL of 2x QuantiTect Probe RT-PCR master Mix, 1 μL of 25 mM MgCl2 (Invitrogen), 4.5μL of RNase-free water, 2 μL each of forward and reverse primer (10μM), 0.25 μL of probe (10μM), and 1 μL of QuantiTect RT Mix. All TaqMan RT-qPCR assays were performed on the thermocycler Rotor Gene Q MDx (Qiagen, Courtaboeuf, France). Amplification was carried out according to the following thermocycling conditions: 50 °C for 30 min for reverse transcription, followed by 95 °C for 15 min and then 45 cycles of 94 °C for 30 s, 55 °C for 30 s and 72°C for 30 s. Negative and positive controls were included in each RT-qPCR assay.

The titer determination of SARS-CoV-2 RNA in number of copies/μL was determined by testing six 10-fold dilutions (i.e. 1.05.10^8^ to 1.05.10^3^ genome copies/mL of a quantitative synthetic RNA from SARS-CoV-2 (BEI Resources). A threshold setting (Ct) of 0.03 was used as the reference threshold for each RT-qPCR assay. The efficiency, slope and correlation coefficient (R^2^) were calculated by the Rotor Gene software. All reactions were carried out as technical duplicates. A cut-off > 36 was defined for negative results and between 32 and 36 for weak positive results (samples with late C_T_ values).

### 4.7. End-point RT-PCR of pol gene of coronaviruses using conserved primers

RNA coronavirus detection was performed by amplifying a 438-bp fragment of the RNA dependent RNA polymerase (pol) gene of coronaviruses using the following degenerated primers: PanCoV pol 15197 (forward): 5’-GGTTGGGAYTAYCCWAARTGTGA, PanCoV pol 15635 (reverse): CCATCRTCMGAHARAATCATCATA designed by (27). The end-point RT-PCR was performed in a two-step RT-PCR with synthesis of cDNA from the extracted total RNA followed by a touch down PCR.

The cDNA was synthesized by reverse transcription of 5μL of RNA extracted from tracheal swabs and faecal samples using 0.25μL of hexanucleotide primers (0.2μG/μL) (Thermo Fisher Scientific, Dardilly, France) and a RT Maxima H Minus cDNA synthesis kit (Invitrogen, Thermo Fisher Scientific, Dardilly, France) according to the manufacturer’s instructions. The cDNA synthesis was performed for 10 min at 25°C, 30 min at 50°C following a final step of 5 min at 85°C. cDNA was stored at < -70°C.

PCR was performed in a final volume of 25 μL containing 3 μL of cDNA, 2.5 μL of 10X PCR Buffer (Invitrogen, Marseille, France), 0.75 μL of MgCl_2_ (50 mM), 1 μL of dNTPs (10 mM) and 0.5 μL of Platinum Taq DNA polymerase (5 U/ml) (Invitrogen, Marseille, France) and 1 μL of forward and reverse primer (20 μM). The PCR was amplified for 2 min at 94°C, with 11 cycles of 30s at 94°C, a 1° touch down decrease of the annealing temperature from 60° to 50°C, 90s at 72°C, then 40 cycles of 30 s at 94°C, 45 s at 50°C and 90 s at 72°C and followed by a final step of extension of 10 min at 72°. Negative and positive controls were included in each RT and PCR assay.

The amplification of the beta-actin gene was performed for each fecal sample with the forward (5’-CGATGAAGATCAAG/ATCATTGC-3’) and reverse (5’-AAGCATTTGCGGTGGAC-3’) primers, with the same methodology to confirm the absence of PCR inhibitors in samples. The amplified products were analyzed by electrophoresis on a 2% agarose gel stained with a SYBR safe solution at a final concentration of 1/10,000 then photographed.

### 4.8. Sequencing of PanCoV amplicons, alignment of sequences and phylogeny

The positive PCR products were sequenced in both directions by Eurofins Genomics (Germany) with the same specific primers used in the PCR.

A dataset of sequences was constituted with 8 sequences from this study (7 pharyngo-tracheal swabs and 1 fecal sample) and 41 referenced sequences including representative sequences of the genus *Alphacoronavirus (n=25), Betacoronavirus (n=20), Deltacoronavirus* (n=4) and *Gammacoronavirus (n=4)*. Multiple alignment of partial pol sequences (positions 14113 to 14536 compared to the reference genome of a *Minacovirus* Strain MW248736) was performed with Mega v10.1.8. A phylogenetic tree was constructed using the Ml method (GTR model). Node robustness was estimated using bootstrap method with 1000 iterations. The consensus sequences of the partial genome of pol gene is accessible in GenBank under accession numbers ON985270 to ON985277.

### 4.9. NGS: Non-specific nanopore sequencing

#### RNA extraction and removal of genomic DNA

Swabs supernatant (150μl) was submitted to RNA extraction using the EZ1 RNA Tissue Mini kit (QIAGEN) following manufacturer’s instructions. Genomic DNA was then depleted from the eluate by incubation with Turbo DNA-free kit (Thermo Fisher Scientific), according to the manufacturer’s instructions.

#### Qualification of the sample by quantification of the SARS-CoV-2 N gene

Quantification of the N gene in the eluate was performed by real-time reverse-transcription (RT-qPCR) using SuperScript™ III Platinum™ One-Step Quantitative RT-PCR System (Invitrogen) as described previously with minor modifications (26). Briefly, a 25 μL reaction contained 5 μL of RNA, 12.5 μL of 2 × reaction buffer, 1 μL of reverse transcriptase/ Taq mixture, 0.4 μL of a 50 mM magnesium sulphate solution (Invitrogen), 1 μl of a 10μM primer, and 0.5l μl of a 10μM probe TxRd-BHQ2. Thermal cycling was performed at 50 °C for 15 min for reverse transcription, followed by 95 °C for 2 min and then 45 cycles of 95 °C for 15 s and 60 °C for 30 s using a Light Cycler 480 (Roche).

#### Ribosomal RNA depletion and real-time reverse-transcription PCR

Eukaryotic rRNA was depleted using the NEBNext rRNA Depletion Kit (Human/Mouse/Rat). After rRNA depletion, cDNA was synthesized from residual total RNA by RT-VILO (Invitrogen) reaction following manufacturer’s instructions. Random amplification or the material was then performed with QuantiTect Whole Transcriptome kit (QIAGEN) according to the manufacturer’s protocol. Amplified DNA was then purified using AMPure XP beads and submitted to Qubit quantification using dsDNA BR Assay Kit and Qubit 3.0 fluorimeter (Invitrogen). Complementary to the un-targeted approach, we also used an adapted version of the published protocol from the ARTIC Network (28) using ARTIC primer scheme version 3, which produces ∼400 bp overlapping amplicons over the SARS-CoV-2 genome.

#### ONT library preparation and MinION sequencing

For maximizing the read length, libraries were not sheared. Sequencing libraries and sequencing reaction were performed according to manufacturer’s instructions with minor adaptations. Briefly, we used the NEBNext Ultra II End Repair/dATailing module (E7546S, NEB, USA) to prepare 1000 ng DNA from each sample. Native barcode adapters NBD04 were ligated in Blunt/TA Ligase Master Mix (M0367S, NEB, USA), and resulting product was AMPure XP beads purified before pooling to produce a 54 μl equimass pool, itself ligated to adapter using Native Barcoding Adapter Mix (BAM). The purified final library was loaded onto an R9.4 flowcell (FLO-MIN106, Oxford Nanopore Technologies, UK), and the run was performed on a MinION Mk1B device (ONT) for 2 hours in order to obtain more than 12 Go of raw data.

#### Genome assembly

Following the MinION sequencing run, raw data were basecalled and reads subsequently demultiplexed using Guppy GPU basecaller / barecoder (Oxford Nanopore Technologies). Raw reads were cleaned using porechop (29) and then mapped against a custom reference of SARS-CoV-2 genome comprising four Chinese and 70 early French sequences using Bowtie2 (30) and minimap2 (31). Finally, consensus genome sequences based on mapped reads (above 83 % coverage with average depth above 100 and a maximum coverage depth over 3000) was generated with bcftools consensus (32).

#### Illumina sequencing

In addition to nanopore sequencing, Illumina fastq files data were also obtained from Illumina sequencing at the National Reference Centre for respiratory viruses (Institut Pasteur Paris). Basecalling and demultiplexing were done on the sequencer using the manufacturer’s software (Illumina). Reads were then trimmed and filtered using Alien trimmer in order to remove adapters and bases under q20 quality score. Mapping was performed on the same reference as mentioned above using Bowtie2 and consensus (majority) genomes were extracted using bcftools and compared to nanopore data. In (rare) case of discordance, higher coverage Illumina data replaced lower coverage nanopore data. After alignment and manual verification, genome sequences were then submitted in GISAID under numbers (EPI_ISL_1392906 & EPI_ISL_10036487-97).

### 4.10. SARS-CoV-2 genomic datasets, genetic and phylogenetic analyses

Three main independent analyses were performed, one using human-derived SARS-CoV-2 genomes, another using mink-derived SARS-CoV-2 genomes and the last one combining all datasets. Firstly, the human SARS-CoV-2-genomes raw dataset was assembled in June 2021 from a collection of all genomes available in GISAID database and filtered on the following parameters: collection date ranging from 01 March 2020 to 30 March 2021, France and human origin, complete genomes, high coverage and low coverage excluded (n=42700). Resulting dataset was aligned using MAFFT (33) and optimized using in-house scripts from sequencing lab of the virology unit, Caen University by removing redundant sequences and similar genomes without losing significant diversity (99,875 % pairwise nucleotide identity cut off, 437 genomes with at least 37 nucleotides differences). This diversity optimized dataset was aligned with the twelve genomes obtained from the French mink farm and analyzed by both Maximum Likelihood (ML from PhyML, SeaView - (34)) and Bayesian phylogenetic methods (Beast - (35)). Both methods used a GTR model of evolution with gamma distribution and invariable sites parameters with the coalescent constant size model as tree prior. Likelihood ratio test and posterior probabilities values were used to estimate node support in ML and Bayesian methods, respectively. The Bayesian phylogenetic analysis was also enriched by using collection dates as priors and an uncorrelated relaxed clock model with lognormal distribution (36). The Markov chain was launched for 100 million iterations on an 18 double cores computer using Beast 1.10.4 suite in order to reach an effective sampling size over 200 for each statistic. The maximum credibility tree was computed from a sampling of 10000 trees and after discarding the first 1000 trees considered as burnin.

Secondly, the mink SARS-CoV-2 genomes raw dataset was assembled using 821 mink SARS-CoV-2 genomes collected from GISAID, with same filtering options previously used for the human SARS-CoV-2 genomes collection and last updated on 5 January 2022. A diversity optimized dataset was generated using the same method as previously described above for the human SARS-CoV-2 dataset. This diversity optimized (SARS-CoV-2 collected in mink - *Neovison vison*) dataset was analyzed by maximum likelihood phylogenetic method with the twelve genomes obtained from the French mink farm. The GTR model of evolution was used with gamma distribution and invariable sites parameters. Likelihood ratio test values were calculated to estimate node support.

Thirdly, additional analyses (SNPs, Amino acid variability) used representatives (n=9) of each main clade identified from previous analyses. Only complete genomes were conserved in this refined dataset and those counting several missing data covering more than 200 contiguous nucleotides in variable loci were discarded. This last dataset was aligned with the twelve genomes sequenced in this study and submitted to ML analyses and variable positions were extracted in order to test the existence and congruence of synapomorphic variations supporting clades and French mink SARS-CoV-2 monophyly.

### 4.11. High throughput sequencing from fecal and swab samples

In addition to the non-specific Nanopore and Illumina sequencing performed on pharyngo-tracheal swabs shown positive for SARS-CoV-2, we undertook high throughput sequencing (HTS) on two pharyngo-tracheal swabs (samples 61-B06-11 and 61-B12-04) and one fecal (sample 61-B23-05) sample, shown positive by conventional RT-PCR for the presence of partial Alphacoronavirus pol gene. The swab and fecal samples were prepared as follows for HTS.

#### Preparation of RNA samples

- Viral RNA extraction was performed for the two swab samples subjected to HTS from a volume of 140 μL using Qiagen Viral RNA mini kit (Qiagen, France) according to the manufacturer’s instructions. Prior to HTS, the two extracted RNA samples were checked for the presence of partial pol gene by the conventional Coronaviruses RT-PCR using the forward and reverse primers PanCoV pol 15197 (F) and PanCoV pol 15635 (R).
- 100 mg of fecal sample were placed in a Virocult tube (Sigma) containing 1 mL of stabilizing buffer, then vigorously vortexed for 30 sec at 7.0 m/s six times in a Fast Prep-24 ™ 5G bead beater (MP Biomedicals Germany Gmbh, Eschwege, France) and centrifuged at 10,000 x g for 5 min. A volume of 210 μL of supernatant was transferred to three individual tubes (i.e. 70μL/tube) for RNA extraction, each filled with 1ml of CTAB buffer (Promega France, Charbonnières les Bains, France). The three tubes were placed in a shaking heat block at 65°C, vigorously vortexed for 1 min and mixed with 40 μL of Proteinase K Solution (Promega France, Charbonnières les Bains, France), before an incubation at 70°C for 10 min and a centrifugation step at 10,000g for 5 min. A volume of 300 μL of clear lysate RNA extraction (i.e. 9 tubes) was performed on the Promega Maxwell RSC instrument with the Maxwell RSC PureFood GMO and Authentication kit according to the manufacturer’s instructions. RNA was pooled in a final volume of 900μL and stored at < -70°C. The extracted fecal RNA sample was shown positive by the conventional Coronaviruses RT-PCR using the forward and reverse primers PanCoV pol 15197 (F) and PanCoV pol 15635 (R).

### Whole genome sequencing and sequence analysis

HTS was performed on the three RNA extracts after a step of rRNA depletion with the rRNA depletion kit (NEB, Evry, France), according to the manufacturer’s recommendations. The RNA library was prepared for each RNA sample tested using Ion Total RNA-Seq kit v2 (Life Technologies, Carlsbad, CA, USA) and then sequenced using Ion Torrent Proton technology. The reads were cleaned with the Trimmomatic 0.36 software, followed by bioinformatics analysis as previously described (37,38) using the GenBank Mink *Caliciviridae* reference sequence MF677852.1 for the two swab RNA samples and the Mink coronavirus 1 reference sequence MN535737.1 for the fecal RNA sample to calculate sub-sampling and final alignment.

Sequences obtained for this study are available in GenBank: Bioproject # PRJNA881217 (Sample 61-B12-04), Bioproject # PRJNA881061 (B-23-05), Biosample: SAMN30886030 (Sample 61-B12-04) and Biosample (B-23-05). The consensus sequence of the full-length genome of the mink *Caliciviridae* is accessible in GenBank under the accession numbers OP485683.

## ACKNOWLEDGEMENTS

We would like to thank all the members of the Anses Nancy Laboratory for rabies and wildlife for their great contributions to this work (sampling and testing every day of week), Christophe Cordevant and Gilles Salvat for their support and Gérald Le Diguerher for his effective help in the last farm. We gratefully acknowledge authors from the originating and sequencing laboratories responsible for obtaining the specimens (Tables S4 and S5).

## FUNDING

This investigation received financial support from the World Health Organization (WHO), through a German fund provided to the WHO R&D Blueprint. The authors alone are responsible for the views expressed in this publication and they do not necessarily represent the views, decisions or policies of WHO. E.S.-L laboratory acknowledges funding from Institut Pasteur, from the INCEPTION programme (Investissements d’Avenir grant ANR-16-CONV-0005), from the NIH PICREID program (Award Number U01AI151758) and from the Labex IBEID (ANR-10-LABX-62-IBEID). S. vdW laboratory acknowledges funding from Institut Pasteur, from Santé publique France, from the Labex IBEID (ANR-10-LABX-62-IBEID) and from the H2020 project 101003589 (RECOVER).

## SUPPORTING INFORMATIONS

**Figure S1. Mink SARS-CoV2 mutations GISAID hcov-19 nextstrain genome Nextclade analysis table**

**Figure S2. SARS-CoV-2 genomes of French mink nested within the 20A clade, together with SARS-CoV-2 from Human sampled at the same period of time**

**Table S1. Mink SARS-CoV2 Nextclade analysis**

**Table S2. Mink SARS-CoV2 genomic variable sites**

**Table S3. Clade and Amino Acid modifications_Nextclade**

**Table S4. GISAID_hcov-19_acknowledgement_table_2022_01_20_13**

**Table S5. GISAID_hcov-19_acknowledgement_table_2022_01_05_10**

## Notes

### Competing Interest Statement

The authors have declared no competing interest.

## REFERENCES

1. Oreshkova N, Molenaar RJ, Vreman S, Harders F, Oude Munnink BB, Hakze-van der Honing RW, et al. SARS-CoV-2 infection in farmed minks, the Netherlands, April and May 2020. Euro Surveill Bull Eur Sur Mal Transm Eur Commun Dis Bull. juin 2020;25(23).

2. Boklund A, Hammer AS, Quaade ML, Rasmussen TB, Lohse L, Strandbygaard B, et al. SARS-CoV-2 in Danish Mink Farms: Course of the Epidemic and a Descriptive Analysis of the Outbreaks in 2020. Anim Open Access J MDPI. 12 janv 2021;11(1):164.

3. Fenollar F, Mediannikov O, Maurin M, Devaux C, Colson P, Levasseur A, et al. Mink, SARS-CoV-2, and the Human-Animal Interface. Front Microbiol. 2021;12:663815.

4. Tada T, Dcosta BM, Zhou H, Vaill A, Kazmierski W, Landau NR. Decreased neutralization of SARS-CoV-2 global variants by therapeutic anti-spike protein monoclonal antibodies. BioRxiv Prepr Serv Biol. 19 févr 2021;2021.02.18.431897.

5. Wu K, Werner AP, Moliva JI, Koch M, Choi A, Stewart-Jones GBE, et al. mRNA-1273 vaccine induces neutralizing antibodies against spike mutants from global SARS-CoV-2 variants. BioRxiv Prepr Serv Biol. 25 janv 2021;2021.01.25.427948.

6. Lu L, Sikkema RS, Velkers FC, Nieuwenhuijse DF, Fischer EAJ, Meijer PA, et al. Adaptation, spread and transmission of SARS-CoV-2 in farmed minks and associated humans in the Netherlands. Nat Commun. 23 nov 2021;12(1):6802.

7. Oude Munnink BB, Sikkema RS, Nieuwenhuijse DF, Molenaar RJ, Munger E, Molenkamp R, et al. Transmission of SARS-CoV-2 on mink farms between humans and mink and back to humans. Science. 8 janv 2021;371(6525):172–7.

8. Rabalski L, Kosinski M, Smura T, Aaltonen K, Kant R, Sironen T, et al. Severe Acute Respiratory Syndrome Coronavirus 2 in Farmed Mink (Neovison vison), Poland. Emerg Infect Dis. sept 2021;27(9):2333–9.

9. Hammer AS, Quaade ML, Rasmussen TB, Fonager J, Rasmussen M, Mundbjerg K, et al. SARS-CoV-2 Transmission between Mink (Neovison vison) and Humans, Denmark. Emerg Infect Dis. févr 2021;27(2):547–51.

10. Molenaar RJ, Vreman S, Hakze-van der Honing RW, Zwart R, de Rond J, Weesendorp E, et al. Clinical and Pathological Findings in SARS-CoV-2 Disease Outbreaks in Farmed Mink (Neovison vison). Vet Pathol. sept 2020;57(5):653–7.

11. Balloux F, Tan C, Swadling L, Richard D, Jenner C, Maini M, et al. The past, current and future epidemiological dynamic of SARS-CoV-2. Oxf Open Immunol. 2022;3(1):iqac003.

12. Shuai L, Zhong G, Yuan Q, Wen Z, Wang C, He X, et al. Replication, pathogenicity, and transmission of SARS-CoV-2 in minks. Natl Sci Rev. mars 2021;8(3):waa291.

13. Krüttgen A, Cornelissen CG, Dreher M, Hornef MW, Imöhl M, Kleines M. Determination of SARS-CoV-2 antibodies with assays from Diasorin, Roche and IDvet. J Virol Methods. janv 2021;287:113978.

14. Mylemans M, Van Honacker E, Nevejan L, Van Den Bremt S, Hofman L, Poels J, et al. Diagnostic and analytical performance evaluation of ten commercial assays for detecting SARS-CoV-2 humoral immune response. J Immunol Methods. juin 2021;493:113043.

15. Vlasova AN, Halpin R, Wang S, Ghedin E, Spiro DJ, Saif LJ. Molecular characterization of a new species in the genus Alphacoronavirus associated with mink epizootic catarrhal gastroenteritis. J Gen Virol. juin 2011;92(Pt 6):1369–79.

16. Wilson DJ, Baldwin TJ, Whitehouse CH, Hullinger G. Causes of mortality in farmed mink in the Intermountain West, North America. J Vet Diagn Investig Off Publ Am Assoc Vet Lab Diagn Inc. juill 2015;27(4):470–5.

17. Scarpa F, Sanna D, Azzena I, Cossu P, Giovanetti M, Benvenuto D, et al. Update on the Phylodynamics of SADS-CoV. Life Basel Switz. 11 août 2021;11(8):820.

18. Su S, Wong G, Shi W, Liu J, Lai ACK, Zhou J, et al. Epidemiology, Genetic Recombination, and Pathogenesis of Coronaviruses. Trends Microbiol. juin 2016;24(6):490–502.

19. Tsoleridis T, Chappell JG, Onianwa O, Marston DA, Fooks AR, Monchatre-Leroy E, et al. Shared Common Ancestry of Rodent Alphacoronaviruses Sampled Globally. Viruses. 30 janv 2019;11(2):E125.

20. Evermann JF, Smith AW, Skilling DE, McKeirnan AJ. Ultrastructure of newly recognized caliciviruses of the dog and mink. Arch Virol. 1983;76(3):257–61.

21. Yang B, Wang F, Zhang S, Xu G, Wen Y, Li J, et al. Complete genome sequence of a mink calicivirus in China. J Virol. déc 2012;86(24):13835.

22. Guo M, Evermann JF, Saif LJ. Detection and molecular characterization of cultivable caliciviruses from clinically normal mink and enteric caliciviruses associated with diarrhea in mink. Arch Virol. 2001;146(3):479–93.

23. Monchatre-Leroy E, Lesellier S, Wasniewski M, Picard-Meyer E, Richomme C, Boué F, et al. Hamster and ferret experimental infection with intranasal low dose of a single strain of SARS-CoV-2. J Gen Virol. mars 2021;102(3).

24. Sailleau C, Dumarest M, Vanhomwegen J, Delaplace M, Caro V, Kwasiborski A, et al. First detection and genome sequencing of SARS-CoV-2 in an infected cat in France. Transbound Emerg Dis. nov 2020;67(6):2324–8.

25. Spada E, Vitale F, Bruno F, Castelli G, Reale S, Perego R, et al. A pre-and during Pandemic Survey of Sars-Cov-2 Infection in Stray Colony and Shelter Cats from a High Endemic Area of Northern Italy. Viruses. 3 avr 2021;13(4):618.

26. Corman VM, Landt O, Kaiser M, Molenkamp R, Meijer A, Chu DK, et al. Detection of 2019 novel coronavirus (2019-nCoV) by real-time RT-PCR. Euro Surveill Bull Eur Sur Mal Transm Eur Commun Dis Bull. 2020;25(3).

27. Gouilh MA, Puechmaille SJ, Gonzalez JP, Teeling E, Kittayapong P, Manuguerra JC. SARS-Coronavirus ancestor’s foot-prints in South-East Asian bat colonies and the refuge theory. Infect Genet Evol J Mol Epidemiol Evol Genet Infect Dis. oct 2011;11(7):1690–702.

28. Tyson JR, James P, Stoddart D, Sparks N, Wickenhagen A, Hall G, et al. Improvements to the ARTIC multiplex PCR method for SARS-CoV-2 genome sequencing using nanopore. BioRxiv Prepr Serv Biol. 4 sept 2020;2020.09.04.283077.

29. Wick RR, Judd LM, Gorrie CL, Holt KE. Completing bacterial genome assemblies with multiplex MinION sequencing. Microb Genomics. oct 2017;3(10):e000132.

30. Langmead B, Salzberg SL. Fast gapped-read alignment with Bowtie 2. Nat Methods. 4 mars 2012;9(4):357–9.

31. Li H. Minimap2: pairwise alignment for nucleotide sequences. Bioinforma Oxf Engl. 15 sept 2018;34(18):3094–100.

32. Li H. A statistical framework for SNP calling, mutation discovery, association mapping and population genetical parameter estimation from sequencing data. Bioinforma Oxf Engl. 1 nov 2011;27(21):2987–93.

33. Katoh K, Standley DM. MAFFT multiple sequence alignment software version 7: improvements in performance and usability. Mol Biol Evol. avr 2013;30(4):772–80.

34. Gouy M, Guindon S, Gascuel O. SeaView version 4: A multiplatform graphical user interface for sequence alignment and phylogenetic tree building. Mol Biol Evol. févr 2010;27(2):221–4.

35. Drummond AJ, Rambaut A. BEAST: Bayesian evolutionary analysis by sampling trees. BMC Evol Biol. 8 nov 2007;7:214.

36. Drummond AJ, Ho SYW, Phillips MJ, Rambaut A. Relaxed phylogenetics and dating with confidence. PLoS Biol. mai 2006;4(5):e88.

37. Briand FX, Schmitz A, Ogor K, Le Prioux A, Guillou-Cloarec C, Guillemoto C, et al. Emerging highly pathogenic H5 avian influenza viruses in France during winter 2015/16: phylogenetic analyses and markers for zoonotic potential. Euro Surveill Bull Eur Sur Mal Transm Eur Commun Dis Bull. 2 mars 2017;22(9):30473.

38. Pellerin M, Hirchaud E, Blanchard Y, Pavio N, Doceul V. Characterization of a Cell Culture System of Persistent Hepatitis E Virus Infection in the Human HepaRG Hepatic Cell Line. Viruses. 4 mars 2021;13(3):406.

